# A DNA-origami NanoTrap for studying the diffusion barriers formed by Phe-Gly-rich nucleoporins

**DOI:** 10.1101/2021.02.07.430177

**Authors:** Qi Shen, Taoran Tian, Qiancheng Xiong, Patrick D. Ellis Fisher, Yong Xiong, Thomas J. Melia, C. Patrick Lusk, Chenxiang Lin

**Affiliations:** Department of Cell Biology, Yale University School of Medicine, New Haven, CT, 06520, USA; Nanobiology Institute, Yale University, West Haven, CT, 06516, USA; Department of Molecular Biophysics and Biochemistry, Yale University, New Haven, CT, 06511, USA

**Keywords:** DNA nanotechnology, DNA origami, nuclear pore complex, size selectivity, diffusion barrier, FG-rich nucleoporins

## Abstract

DNA nanotechnology provides a versatile and powerful tool to dissect the structure-function relationship of biomolecular machines like the nuclear pore complex (NPC), an enormous protein assembly that controls molecular traffic between the nucleus and cytoplasm. To understand how the intrinsically disordered, Phe-Gly-rich nucleoporins (FG-nups) within the NPC’s central transport channel impede the diffusion of macromolecules, we built a DNA-origami NanoTrap. The NanoTrap comprises precisely arranged FG-nups in an NPC-like channel, which sits on a baseplate that captures macromolecules that pass through the FG network. Using this biomimetic construct, we determined that the FG-motif type, grafting density and spatial arrangement are critical determinants of an effective diffusion barrier. Further, we observe that diffusion barriers formed with cohesive FG-interactions dominate in mixed-FG-nup scenarios. Our DNA-origami platform thus sheds light on how NPCs sieve inert macromolecules and will provide a valuable tool for studying nuclear transport.

## INTRODUCTION

Nuclear pore complexes (NPCs) reside in the nuclear envelope where they control the bidirectional exchange of molecules between the nucleus and cytoplasm.^1, 2^ An individual NPC is composed of ~30 proteins (nucleoporins or nups) that build an 8-fold radially symmetric scaffold that houses a 40–50 nm wide central channel.^2, 3^ The channel is filled with intrinsically disordered nups rich in phenylalanine (F) and glycine (G) amino acids in repetitive motifs.^2–4^ These “FG-nups” control the passage of molecules through both passive and active mechanisms that ultimately serve to ensure proper nuclear-cytoplasmic compartmentalization.^5^ First, they establish a diffusion barrier that restricts the passive diffusion of macromolecules larger than ~40 kD.^6–8^ Second, they provide binding sites for nuclear transport receptors (NTRs),^9^ that rapidly ferry cargo molecules through the NPC, with energy and directionality contributed by the Ran GTPase.^10^ A mechanistic understanding of the NPC’s gatekeeping function is not only essential to the study of cellular compartmentalization, but also provides design inspirations for synthetic macromolecule-sorting machines.

Work over the last few decades has sought to conceptualize the FG-nups as forming higher-order assemblies bearing a physicochemical resemblance to “hydrogels”,^11–13^ “polymer brushes”^14–18^ and most recently, liquids.^19^ Such models differ based largely on the relative importance attributed to cohesive interactions between FG-nups, which have been observed *in vitro*^8, 12^ but may be mitigated *in vivo* due to non-specific competition.^18^ Some *in silico* models suggest that cohesive interactions among the FG-network exist in a regime that allows for reversible condensation, a putative key element of the gating mechanism.^20^ The lack of consensus here is also driven by the observations that cohesive and non-cohesive nups occupy different regions of the central channel, which may suggest discrete local environments that may or may not impact the entire collective.^21–23^ Exploring the potential function of FG-nups requires more elaborate NPC mimics, which so far have been limited to lining nanometer-sized artificial channels with a single type of FG-nup.^24, 25^ As impressive as the current NPC mimics are, it remains challenging to delineate the contributions of precise numbers of different individual FG-nups, in particular their amino-acid compositions and positioning within the nuclear pore, to the overall permeability and selectivity of the FG-nup collective that makes the NPC function.

To reveal the structure-function relationship of the NPC’s central channel, namely how FG-nup configurations affect the formation of a size-selective diffusion barrier, we have built a DNA-origami-based biomimetic assembly that we term NanoTrap. NanoTrap allows the precise positioning of FG-nups with diverse FG-motifs (e.g., FxFG, GLFG, FG, SAFG, and PSFG)^5, 26^ within a DNA channel that gate the entry of proteins into a sealed chamber (**Figure 1**). This work thus takes advantage of the well-defined shape, programmable assembly and the chemically addressable surface of DNA-origami structures,^27–30^ while building on our recently reported NuPOD (NucleoPorins Organized on DNA) platform.^31^ The latter (and a similar platform)^32^ enabled the study of the morphology and dynamics of FG-nup collectives anchored on DNA-origami channels.^31^ Similar to the prior study, here we systematically vary the organization of two representative yeast FG-nups, Nsp1 (FxFG-rich) and Nup100 (GLFG-rich), to study how cohesiveness, density, and spatial organization of these FG-nups impact their selective sieving behaviors in a confined NPC-channel-like space. However, unlike the first-generation NuPODs, the NanoTrap with a macromolecule-trapping baseplate allows the direct assessment of the ability of the FG-nup collectives to filter biomolecules of different molecular weights (i.e., size selectivity), thereby establishing a functional assay for FG-nups in a well-controlled, nanopore-like environment. We expect this adaptable platform to help better define the properties of the NPC central channel, and ultimately contribute to elucidating the underlying mechanisms of nuclear transport.

**Figure 1.**
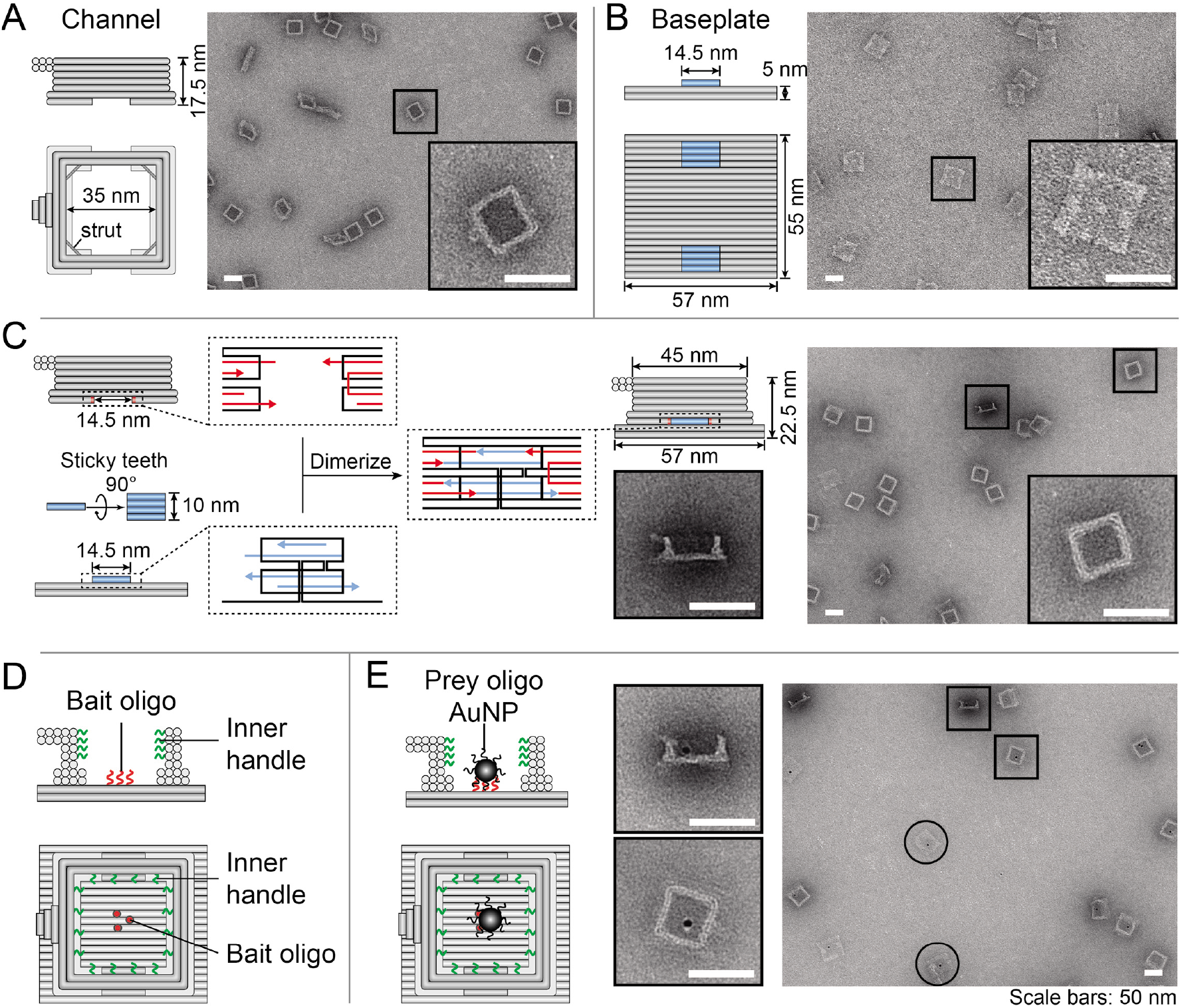
A DNA-origami NanoTrap built from two pre-assembled parts, a channel and a baseplate. (A) Cartoon models and negative-stain EM images of the channel. (B) Cartoon models and negative-stain EM images of the baseplate. (C) Two teeth (blue) with sticky ends (blue arrows) mediate the baseplate attachment onto the channel, which contains two cavities with complementary sticky ends (red arrows), resulting in the formation of a NanoTrap (cartoon model and negative-stain EM images shown on the right). (D) Up to four layers of DNA handles (12 handles per layer) protrude from the channel wall (green curls), serving as anchor points for anti-handle-conjugated proteins. The baseplate displays three “bait” oligonucleotides (red curls/dots) to capture nanoscale objects carrying “prey” oligonucleotides. (E) Cartoon models and EM images showing the immobilization of prey-oligo-modified AuNPs (dark spots) inside the NanoTrap. Circles indicate standalone baseplates or “flipped over” Nanotraps. Scale bars = 50 nm. See also **Figure S1**, **Figure S2**, **Figure S3**, and **Figure S4**.

## RESULTS

### A DNA-origami NanoTrap mimics key features of the NPC central channel

To engineer a molecular environment that recapitulates the essential structural and biochemical properties of the NPC central channel, we designed a NanoTrap consisting of two DNA-origami components: a channel and a baseplate (**Figure 1** and **S1**). The cuboid-shaped channel has an inner width of 35 nm and a height of 17.5 nm (**Figure 1A**), forming the entryway of the NanoTrap. On the interior wall of the channel extend DNA oligonucleotides (handles) for hybridization with complementary DNA (anti-handles) that are conjugated to FG-nups. Each DNA channel is equipped with up to 48 handles, distributed in four layers (12 handles/layer) with radial symmetry (**Figure 1D**). The handle sequences are independently tunable, allowing us to decorate designated positions (layers) of the channel with selected FG-nups to build different NanoTraps, which we denote according to the protein type, stoichiometry, and anchor position (e.g., Nup10024Nsp124 for a channel with top two layers of handles occupied by a total of 24 copies of Nup100 and the bottom two layers by 24 copies of Nsp1).

To detect macromolecules passing through such an NPC mimic, we sealed the channel bottom by building a DNA-origami baseplate (57 nm×55 nm×5 nm, **Figure 1B**) with two raised stacking interfaces or ‘teeth’ (**Figure 1B**, blue bars). Each tooth comprises four parallel DNA helices, with a shape and sticky-end sequences complementary to a corresponding cavity on the bottom of the DNA channel that allows assembly of the complete NanoTrap (**Figure 1C**). The baseplate displays three single-stranded DNA extensions (termed ‘bait’ oligonucleotides) to immobilize entering molecules modified by a ‘prey’ oligonucleotide. As a result, macromolecules may only enter the NanoTrap through the entryway, subject to filtration by the FG-nups, and those that do cross the FG-nup barrier to reach the baseplate will be captured for easy detection by fluorescence-based assays.

We verified the proper formation of the DNA channel, baseplate, and the NanoTrap using well established DNA-origami assembly, purification, and characterization methods.^31, 33^ Gel electrophoresis (**Figure S2**) and negative-stain electron microscopy (EM) (**Figure 1**) clearly showed the correctly formed DNA-origami structures with expected shapes and dimensions. Importantly, stable NanoTraps formed with high efficiency (~80% dimerization yield) *via* the straight teeth and cavities, in contrast to an antecedent design of a cylindrical channel with curved stacking interfaces that led to inefficient dimerization (**Figure S3**). We further examined the chamber’s molecular trapping ability using gold nanoparticles (AuNPs) to mimic entering nanoscale entities. These 5 nm AuNPs were first conjugated with prey oligonucleotides (**Figure S4**) and then incubated with empty NanoTraps (without FG-nup grafting) that display baits on their baseplates. Negative-stain EM confirmed that ~85% of NanoTraps had AuNP attachment, demonstrating good molecular accessibility to baseplates and a high capture rate of prey-modified molecules (**Figure 1E**). The AuNP-free NanoTraps may be caused by baseplates missing their bait oligonucleotides during DNA-origami folding and/or a small portion of AuNPs losing their prey oligonucleotides after DNA conjugation.

The NPC’s central channel comprises a variety of nucleoporins with diverse FG-repeats that also differ in their propensity to interact with themselves and other FG-repeats.^8, 34^ Previous studies have shown that cohesive FG-nups (e.g., Nup98, the human orthologue of Nup100) are essential to the NPC’s function as a permeability barrier.^12, 35^ The DNA-origami-based NanoTrap enables a bottom-up approach to test the role of different FG-repeats in forming a size-selective barrier, owing to the complexity-reduced *in vitro* system and the exquisite control over FG-nup placement. To demonstrate the concept, we tested two common FG-motifs, GLFG, and FxFG. The more cohesive GLFG-rich domain^36, 37^ of Nup100 (amino acids 2–610) and the less cohesive FxFG-rich domain^37, 38^ of Nsp1 (amino acids 2–603) were produced in *E.coli* and affinity purified with an MBP-SUMO tag (to improve stability and solubility) and a SNAP tag (for conjugation with benzylguanine labeled DNA anti-handles). After conjugation with the DNA anti-handle, the MBP-FG-nup-domain-DNA conjugates (FG-nups hereafter for simplicity) were purified from free MBP-FG-nups by size-exclusion chromatography: the conjugation yield was above 90% as verified by SDS-polyacrylamide gel electrophoresis (SDS-PAGE; **Figure 2A** and **S5**). The FG-nup-gated NanoTraps were assembled by incubating purified, undecorated NanoTraps with anti-handle-conjugated nucleoporins. With all 48 handles occupied, the NanoTrap reaches a grafting density of ~3 FG-nups per 100 nm^2^ and an FG-repeat concentration of up to 150 mM, similar to those in the NPC central channel.^15, 39, 40^ The final assemblies were characterized by SDS-agarose electrophoresis and negative-stain EM (**Figure 2B, S6** and **S7**).

**Figure 2.**
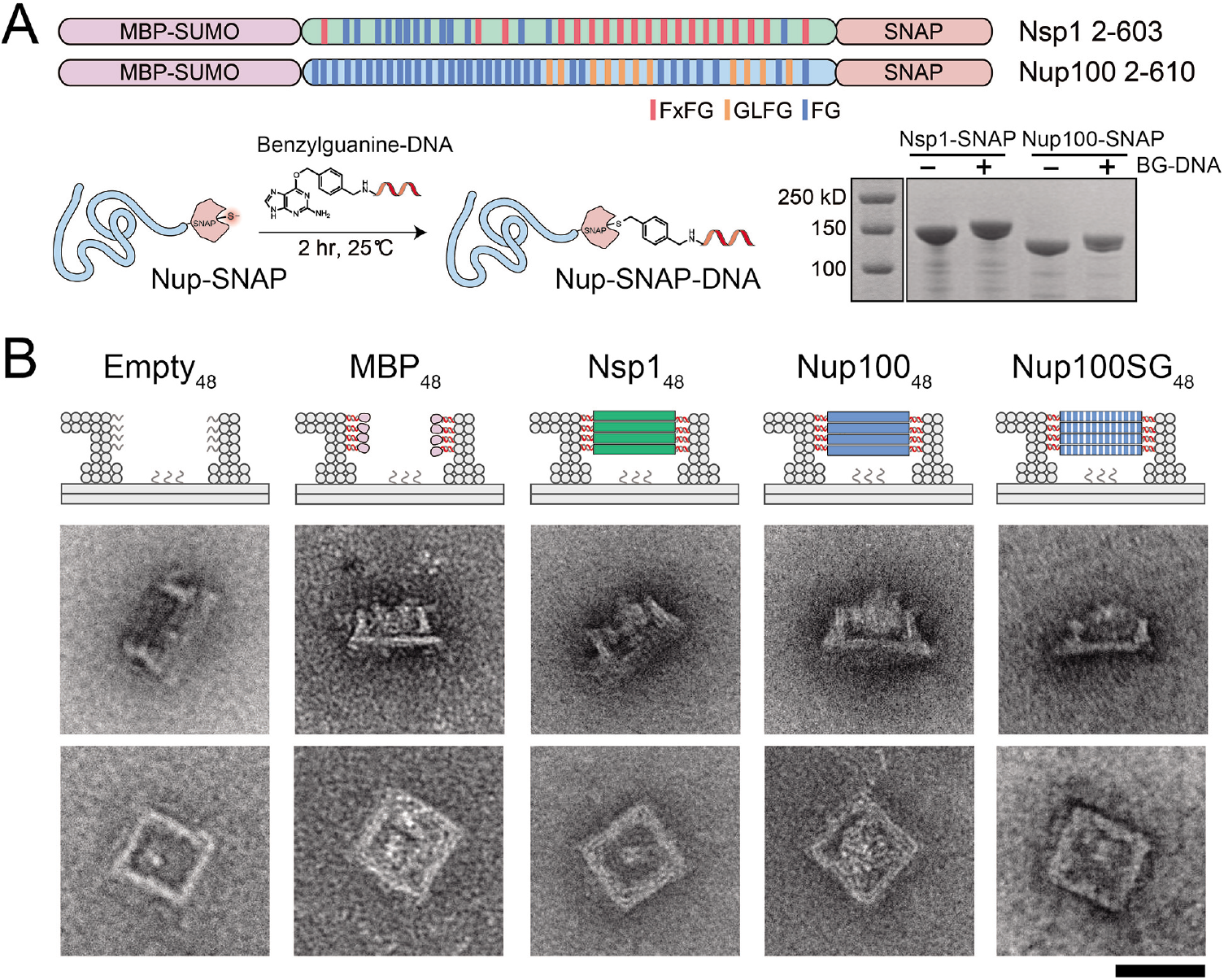
Assembly and characterization of nucleoporin-gated NanoTraps. (A) Two central channel FG-nup domains of yeast origin, Nsp1 (FxFG-rich) and Nup100 (GLFG-rich), were cloned and expressed in *E. coli* as MBP-SUMO-nup-SNAP fusions. Such SNAP-tag bearing nucleoporins were conjugated with benzylguanine modified DNA oligonucleotides (BG-DNA) and verified by SDS-PAGE. (B) Cartoon models (top) and negative-stain EM images (bottom) of an ungated (empty) and various protein-gated NanoTraps. Scale bar = 50 nm. See also **Figure S5**, **Figure S6** and **Figure S7.**

### Type of FG-repeat affects the strength of diffusion barrier

The NanoTrap design facilitates convenient ensemble measurement of the amount of macromolecules that cross the FG-nup barrier, thereby enabling systematic testing and unambiguous ranking of barrier strengths. We started by testing the diffusion barriers formed by the different FG-nups by incubating the NanoTraps (3 nM) with a GFP-tagged prey-oligo-conjugated protein (GFP-SNAP-prey, 53 kD, 1 μM), which would be immobilized on the baseplate upon entering the trap (**Figure 3A**). A given barrier’s permeability was determined by separating the reaction mixture by gel electrophoresis and normalizing the NanoTrap’s GFP fluorescence (from the trapped molecules) against its ethidium bromide fluorescence. We first ensured that the GFP-SNAP-prey conjugate had unfettered access to the baseplate when no FG-nups decorated the channel. Indeed, after incubation with the GFP-SNAP-prey conjugate (**Figure S8**), the undecorated trap exhibited clear GFP fluorescence (**Figure 3B**), which we set as the maximal penetration (100%) for subsequent quantification of barrier permeability. For example, we found that 48 copies of the maltose-binding proteins (MBP, ~40 kD) did not hinder access of the 53 kD protein, as the MBP_48_ channel allowed nearly as much (~95%) GFP-binding on the baseplate as an empty channel did (**Figure 3B**). In contrast, the NanoTrap with 48 copies of Nsp1 gating its entryway (Nsp1_48_) halved the penetration of the 53 kD protein, suggesting that it imposed a barrier to this molecule. Interestingly, the Nup100_48_-gated trap was virtually impermeable with only ~7% penetration of the GFP-SNAP-prey reporter. These data suggest that compared with the well-folded MBPs, the intrinsically disordered FG-nups can impose diffusion barriers to at least a 53 kD protein, albeit with different characteristics. Indeed, the relative permeability of the NanoTraps could be predicted from our studies of the first-gen NuPODs, which showed that while MBP tended to occupy space near the channel wall, Nsp1 could sample volume both inside and outside of the channel and Nup100 formed a plug-like condensate.^31^ Thus, these morphological differences correlate with the permeability of the NanoTraps and suggest that the propensity to form a cohesive network contributes to barrier strength.

**Figure 3.**
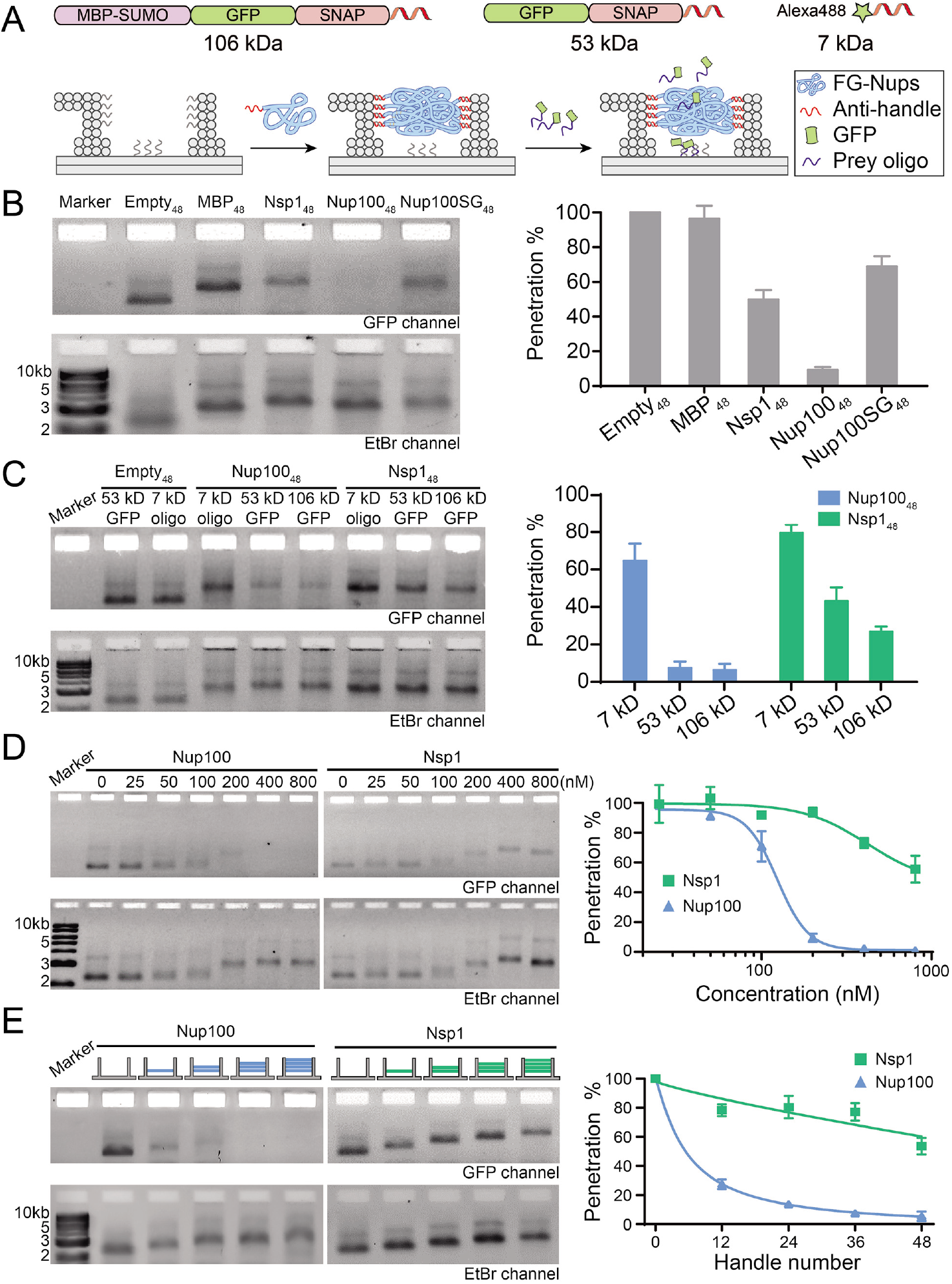
The influence of FG-nup type, density and geometric distribution on the barrier permeability. (A) Schematic diagrams showing the permeability assay using an FG-nup-gated NanoTrap and fluorescently tagged macromolecules of different sizes (106 kD MBP-GFP-SNAP-prey, 53 kD GFP-SNAP-prey, and 7 kD Alexa488-prey). (B) The barrier strengths of different gating proteins (48 copies per NanoTrap) against the 53 kD GFP-SNAP-prey. (C) Size-selective diffusion barriers formed by 48 copies of nucleoporins (Nup100 or Nsp1) within a NanoTrap. (D) Permeability of NanoTraps (4 nM, 48×handles per trap) formed at different nucleoporin (Nup100 or Nsp1) concentrations (0–800 nM), tested against the 53 kD GFP-SNAP-prey. Fitted curves are guides to the eye. (E) Permeability of NanoTraps containing 12–48 copies of Nup100 or Nsp1 tested against the 53 kD GFP-SNAP-prey. The exact nup arrangement is shown by the schematic drawing at the top of each lane (blue: Nup100, green: Nsp1, 12 nups/layer). Fitted curves are guides to the eye. Statistical data are plotted to show mean ± standard error of the mean (SEM) from three trials. See also **Figure S8**.

Consistent with cohesive interactions contributing to barrier strength, our results showed that the impermeability of the Nup100-NanoTraps could be overcome by disrupting cohesive interactions mediated by the GLFG repeats. We generated a NanoTrap gated by 48 copies of a Nup100 variant where every phenylalanine (F) in the 44 FG repeats was mutated to serine (S) that is known to reduce the cohesiveness of FG-repeats^12^. Interestingly, while these mutations led to an increase in permeability of the Nup100SG_48_-NanoTraps with ~70% penetration, they were able to form a diffusion barrier with similar permeability as Nsp1_48_ (**Figure 3B**). Thus, while GLFG-mediated cohesive interactions impact the relative strength of the diffusion barrier, FG-repeats *per se* are not required to impede the passage of the GFP-SNAP-prey.

### GLFG and FxFG nups have unique size-selective filtering properties

To gain more insight into the diffusion barriers established by Nsp1 and Nup100, we next tested a series of reporters that ranged from 7 kD to 106 kD (**Figure 3A** and **3C**). Both Nsp1_48_ and Nup100_48_ imposed only slight impedance to the passage of the 7 kD DNA molecule (~70–80% penetration); importantly, these data also established that the FG-nups minimally interfered with binding of the prey oligonucleotide to the baseplate. Interestingly, while Nup100-NanoTraps were essentially impermeable to both the 53 and 106 kD preys, Nsp1-NanoTraps allowed passage of both in a manner that correlated with their molecular weights (the 53 kD and 106 kD reporters penetrated to ~50% and ~30%, respectively). Thus, while Nup100 appears to establish a diffusion barrier with a more stringent molecular weight cutoff (<53 kD), suggesting the formation of a sieve-like hydrogel, Nsp1’s permeability to macromolecules is more consistent with an entropy-driven barrier, in that molecules with increasing molecular weight are met with more resistance. These results suggest the NanoTrap’s size selectivity, either “soft” or “hard”, is heavily influenced by the properties of the constituent FG-nup, making the programmable FG-nup-gated NanoTraps an ideal *in vitro* platform for studying the molecular underpinnings of the NPC’s size selectivity.

### Densely grafted nucleoporins under spatial confinement lead to strong barriers

In the experiments described above, we crowded 48 copies of FG-nups into the NanoTrap to match the FG-repeat concentration in a natural NPC. To understand how FG-repeat density could influence barrier properties, we performed two sets of experiments. First, we titrated 4 nM of NanoTraps bearing 48-handles with increasing concentrations (0–800 nM) of Nsp1 or Nup100 as a means to gradually increase the average FG-nup density within the NanoTraps until saturation. The resulting FG-nup-gated NanoTraps were subjected to the permeability assay (**Figure 3D**). Increasing FG-nup concentration led to slower mobility of the NanoTraps in the gels as well as decreased penetration of the 53 kD GFP-SNAP-prey. Notably, both the NanoTrap mobility and permeability of the Nup100-gated NanoTraps remained mostly unchanged when the Nup100 concentration was below 50 nM (i.e., on average ≤12.5 Nup100 per trap), but dropped dramatically when the Nup100 concentration was raised from 100 nM to 200 nM (i.e., on average ≥25 Nup100 per trap). Further increasing nucleoporin concentration produced diminishing improvements to the barrier strength. This sharp transition suggested a possible concentration-dependent, cooperative association among the FG-nups within a nanopore — only when FG-repeats reach a minimal density does a functional barrier form. At every FG-nup concentration tested, Nsp1 formed a weaker barrier than Nup100, further supporting a cohesiveness-dependent permeability barrier.

Second, we varied the number of handles in the NanoTrap to deterministically control the FG-nup organization (**Figure 3E**). We thus derived eight versions of NanoTraps, each with 12, 24, 36, or 48 copies (i.e., 1–4 layers) of Nup100 or Nsp1 gating the entryway of a NanoTrap. Consistent with the data presented above, Nsp1 alone could not form a sufficient barrier to block the 53 kD GFP-SNAP-prey from entering the NanoTrap, except at the highest FG-nup densities tested where a moderate impact was observed (**Figure 3B–D**). In contrast, merely one layer (12 copies) of Nup100 placed near the bottom of the DNA channel (Empty36Nup10012) reduced GFP-SNAP-prey penetration to ~30%. Additional Nup100 produced moderate enhancement of the barrier strength, with a near-complete rejection of the GFP-SNAP-prey with 24 or 36 copies of Nup100. This result is in qualitative agreement with the sharp transition of permeability observed when titrating the Nup100-to-NanoTrap ratio (**Figure 3D**).

Interestingly, 12× Nup100 formed a better diffusion barrier when grafted in one layer as opposed to when it was randomly distributed across all 4 layers as would be predicted to be the case in the Nup-to-NanoTrap titration experiments. These data suggest that, additional confinement endowed by spatial positioning may also play a role in the formation of a diffusion barrier. To test this hypothesis, we attached the same copy numbers of nucleoporins at different axial positions (i.e., layers) along the entryway of the NanoTrap. As shown in **Figure 4A**, providing there were at least 24 copies of Nup100, changing their axial positioning did not lead to an observable impact on reporter penetration. However, we observed a significant permeability difference when only 12 copies of Nup100 were grafted at either the top or bottom of the chamber, with (Nup10012Empty36) showing ~70% penetration compared to Empty36Nup10012 with only ~30% penetration. We interpret these data in a model in which Nup100 near the channel opening is less likely to form a tightly-knit network because of its ability to sample larger solvent volume. Thus, our data highlight the importance of geometric constraints in forming an FG-nup-based diffusion barrier, at least in one that relies on cohesive interactions, which are likely favored under confinement.

**Figure 4.**
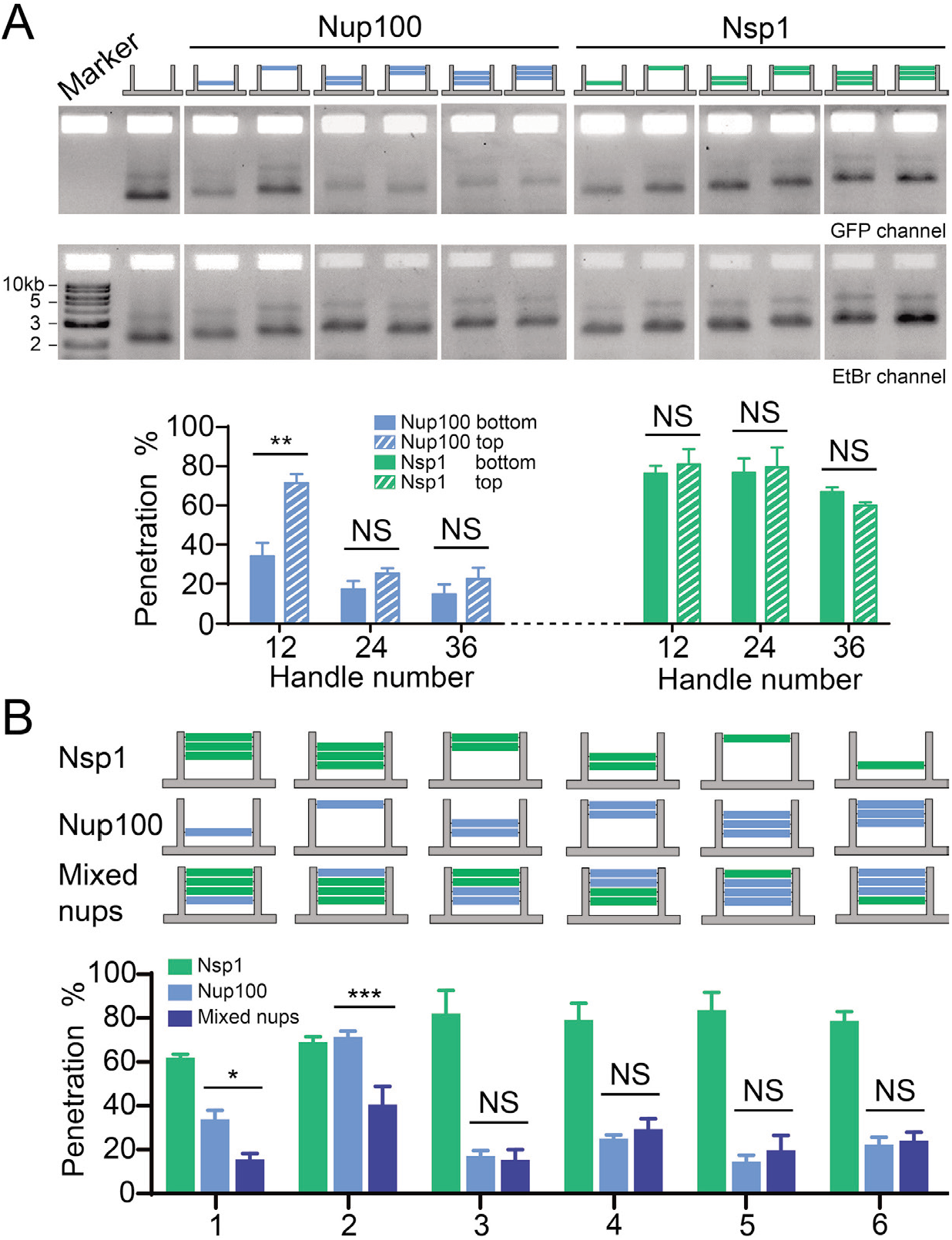
Different nucleoporin arrangements affect barrier permeability. (A) Permeability of NanoTraps with the FG-nups located near the entrance (top) or the baseplate (bottom) of the NanoTrap, tested against the 53kDa GFP-SNAP-prey. The exact nup arrangement is shown by the schematic drawing at the top of each lane (blue: Nup100, green: Nsp1, 12 nups/layer). Statistical data are plotted to show mean ± SEM. Statistical significance was determined by a two-tailed Student’s t-test; n=3; NS: not significant (*P*≥0.05); **: *P*<0.01. (B) Permeability of NanoTraps containing single or mixed types of FG-nups, tested against the 53kDa GFP-SNAP-prey. The exact nup arrangements are shown by the schematic drawings at the top of each group of bars (blue: Nup100, green: Nsp1, 12 nups/layer). Data are plotted to show mean ± SEM. Difference between mixed-nup and Nup100-NanoTraps was analyzed by two-way ANOVA and Tukey’s multiple comparison; n=3; NS: not significant (*P*≥0.05); *: *P*<0.05; ***:*P*<0.001.

### The more cohesive Nup100 dictates the barrier strengths of mixed FG-nups

The formation of distinct diffusion barriers by the two different FG-nups raises the question of how, or whether, they impact each other within the NPC-like nanopore confinement. Addressing this question will shed light on how the physiological *in vivo* NPC diffusion barrier is ultimately formed from the collective of a dozen FG-nups with unique biophysical characteristics. To begin to explore how combinations of distinct FG-nups might alter the properties of a diffusion barrier, we anchored both the GLFG-dominated Nup100 and FxFG-dominated Nsp1 into a single NanoTrap. In total, we designed 6 different NanoTrap handle arrangements to facilitate the attachment of mixed FG-nups, as illustrated in **Figure 4B**. Each handle arrangement has three subtypes: one containing Nup100 and Nsp1, one with Nup100 only, and one with Nsp1 only.

Using the SDS-agarose gel assay to detect the penetration efficiency of the 53 kD GFP-SNAP-prey, we observed that in scenarios in which there were 24 copies of Nup100, the resulting cohesive network dominated and was not impacted by the introduction of even 24 copies of Nsp1. Interestingly, however, in cases with only 12 copies of Nup100, the introduction of Nsp1 strengthened the barrier (**Figure 4B**). This effect was most significant when Nup100 was grafted at the entrance to the channel, where Nup10012Empty36 itself did not form an efficient barrier (**Figure 4A**). These results suggest that Nsp1 might promote cohesive interactions of relatively low copy-number networks of Nup100 or that Nup100 promotes the confinement of Nsp1 in a way that establishes a stronger barrier.

## DISCUSSION

The generation of an *in vitro* mimic capable of recapitulating the NPC’s complexity remains a significant challenge for synthetic biology. The NanoTrap system presented here has advanced our NuPOD platform to extend its utility beyond a morphological characterization of FG-nups confined to a nanochannel, to one that is now capable of assessing a key *in vivo* function of the FG-nups: the generation of a diffusion barrier. We draw the following conclusions from this work: First, cohesive FG-nups can form a much stronger barrier to molecular passage than their less-cohesive siblings. Indeed, as few as 12 copies of Nup100, provided they are confined within the channel walls, can form a diffusion barrier to macromolecules as small as 50 kD. Consistent with the notion that the cohesive properties of the GLFG repeats are responsible for this function, the cohesivity-ablating Nup100SG mutant fails to establish such a barrier.^7, 12^ These data are thus congruent with many other studies that have also compellingly established that the cohesivity of nups is a central feature of an effective diffusion barrier, at least *in vitro*.^8, 12, 35^ However, it remains to be established whether the abrupt molecular weight cutoff that we (**Figure 3C**) and others^12^ have observed with Nup100 ultimately reflects the permeability properties of NPCs *in vivo*. Indeed, recent work suggests that the diffusion barrier may be “soft” and permeable to even very large macromolecules.^6, 41^ Our recent work also showed that the tip of the megadalton-sized HIV capsid can at least insert into the NPC-mimicking NuPOD.^42^

The permeability properties of Nsp1 that we measured in the NanoTraps seem to be more in line with a soft-diffusion barrier type mechanism — one with a continuum of penetration rates that negatively correlates with increasing molecular weights of entering molecules. These observations are consistent with entropic barrier models where the dynamic nature of unstructured and flexible FG-nup filaments effectively occlude the channel.^6, 43^ However, a central challenge in the field is to reconcile the abundance of data surrounding the importance of cohesive interactions in controlling the function of the NPC with such an entropic barrier model of the FG-nups. For example, it is potentially interesting that the cohesive and non-cohesive nups seem to populate distinct parts of the central channel,^8, 21–23, 37^ but whether this relative positioning functionally matters remains elusive. Similarly, it remains uncertain how cohesive and non-cohesive nups interact to mitigate or enhance their individual cohesive properties. Our system provides an ideal platform for assessing these questions within the highly controlled environment afforded by the DNA NanoTraps. While we are just beginning these investigations, some interesting themes are emerging. First, a cohesive network of GLFG-nups will dominate over its non-cohesive siblings when at sufficiently high concentrations. These data suggest that non-cohesive interactions are inefficient to weaken a cohesive network. Consistent with this, and secondly, when cohesive nups are at low concentrations, the addition of a non-cohesive network can actually increase the strength of the barrier. Thus, collectively, these data support a model in which cohesive networks would dominate within the native NPC.

Furthermore, our work supports the concept that confinement of FG-nups within a channel impacts their collective properties and favors their condensation.^31^ The observation that best exemplifies this is that 12 copies of Nup100 can only form a barrier when grafted deep within the NanoTrap and not near its entrance, where the nucleoporins are exposed to the large surrounding volume. To reconcile these observations of an *in vitro* cohesive “hard” barrier with the observations of an *in vivo* “soft” barrier, future experiments will need to explore how other nucleoporins as well as factors like NTRs or non-specific competitors ultimately serve to weaken cohesive interactions, as has been suggested.^44^

Beyond providing an adaptable framework for studying nuclear transport mechanisms, the NuPOD/NanoTrap systems can be viewed as prototypes of macromolecule sorting devices with tunable size-filtration behaviors. The impact of the gating biopolymers’ cohesivity and positioning on the NanoTrap’s permeability provide valuable design references for future engineering of DNA nanopores with increasing structural and functional diversity. Future development in this direction, in conjunction with the fast-evolving technologies that shape and perforate membranes with DNA nanostructures,^45–47^ may usher in a range of applications in biotechnology such as sensing viral pathogens and synthetic biology such as building artificial nuclei.

## EXPERIMENTAL PROCEDURES

### Resource Availability

#### Lead Contact

Further information and requests for resources and reagents should be directed to the Lead Contact, Dr. Chenxiang Lin.

#### Materials Availability

Constructs are available either in a public repository or via requests to the corresponding authors.

#### Data and Code availability

All data generated or analyzed during this study are included in this article and its supplementary information files. No unpublished custom code, software, or algorithm was used in this work.

### General

All chemicals were purchased from commercial sources and used without further purification unless otherwise stated. All DNA oligonucleotides were purchased from Integrated DNA Technologies (IDT).

### Cloning and Expression

The coding sequences for amino acids 2–603 of Nsp1 and amino acids 2–610 of Nup100 from *S. cerevisiae* were cloned into 10×His-MBP-SUMO-nup-SNAP constructs (**Figure 2A**) via a pET-28a-derived vector (Novagen), and expressed in *E. coli* strain BL21-Gold(DE3). The coding sequences of GFP were cloned into the same vector with or without the MBP-SUMO tag (**Figure 3A**). The plasmids were transformed into *E. coli* strain BL21-Gold(DE3) competent cells via heat shock. For expression, transformed bacteria were cultured in Luria Broth media with kanamycin (50 μg/mL) at 37°C while shaking at 220 rpm for 4–6 hr, until OD_600_ reached ~0.8. IPTG (1 mM) was then added to induce protein expression for 4–5 hr at 25°C before cell collection by centrifugation. Cell pellets were stored at −80°C until use.

### Proteins purification

The cell pellet was thawed and resuspended in lysis buffer (1×PBS containing 150 mM NaCl, pH 7.4, 0.5 mM TCEP, 0.1 mM PMSF, 1 × Roche complete protease inhibitors), and lysed in a cell disruptor. Whole-cell lysates were spun at 35k rpm for 45 min in a Type 45 Ti rotor (Beckman Coulter), and the supernatant was decanted and filtered through a 0.45 μm cellulose acetate membrane. The resulting filtered lysate was applied to a 5 mL HisTrap column (GE Healthcare) on an ÄKTA FPLC system (GE Healthcare) at a 1 mL/min flow rate. The column was washed with wash buffer (1×PBS, 0.1% Tween 20, 25 mM imidazole) and eluted on a gradient of elution buffer (1×PBS, 0.1% Tween 20, 25–500 mM imidazole). Protein concentration was determined by Nanodrop (Thermo Fisher Scientific). Samples were flash-frozen in liquid nitrogen and stored at −80°C until use.

### Benzylguanine (BG)-DNA preparation

DNA anti-handles (5’-labeled amino-DNA oligonucleotides) were resuspended in deionized H2O at 2 mM. BG-GLA-NHS (New England BioLabs) was dissolved in DMSO at 20 mM. DNA anti-handles were then mixed with BG-GLA-NHS in a 1:3 volumetric ratio in 70 mM HEPES buffer (pH 8.5) and incubated at room temperature (r.t.) for 1 hour. The BG-DNA product was then purified from excess BG-GLA-NHS by ethanol precipitation. Dried BG-DNA pellets were stored at −20°C until use.

### Protein-DNA conjugation and purification

BG-DNA pellets were resuspended in deionized H2O and mixed with purified nups in 1×PBS buffer to reach a final concentration of 40 μM BG-DNA and 20 μM nup (2:1 molar ratio). This reaction mixture was incubated at 25°C for 2 hours. Excess DNA was removed from conjugated proteins using size exclusion chromatography on a Superdex200 10/300 column (GE Healthcare) in a 25 mM Tris buffer (pH 8) containing 75 mM NaCl. Conjugation efficiency was verified by SDS-PAGE (see below).

### SDS-polyacrylamide gel electrophoresis (SDS-PAGE)

All SDS-PAGE gels contained 8% acrylamide bis-tris (Bio-Rad, pH 6.5). Samples were boiled in 1× Laemmli sample buffer at 90°C for 5 mins before loading to the gels. The gels were run for 40 min at 25 V/cm in MOPS-SDS buffer (50 mM Tris, 50 mM MOPS, 1 mM EDTA, 0.1% SDS, pH 6.5). Gels were stained with Coomassie Blue or SYPRO Red (Thermo Fisher Scientific).

### DNA-origami design and assembly

Channel and baseplate were designed in caDNAno^48^ (caDNAno.org), with bait and handle extending from the 3’ end of staple strands at positions indicated in **Figure 1D** and **S1**. The extension sequences are 5’-AAATTATCTACCACAACTCAC-3’ (inner handle a), 5’-CTGATGATATTGATTGAAATG-3’ (inner handle b), and 5’-CTTAAGCGATACGGGAATATG-3’ (bait). The DNA-origami structures were assembled from an M13mp18 bacteriophage-derived circular ssDNA strand (8064 nt) and staple oligonucleotides (see **Figure S1**). The assembly was carried out using a 36 hr 85°C–25°C annealing gradient in 1×TE buffer (5 mM Tris-HCl, 1 mM EDTA, pH 8.0) supplemented with 15 mM MgCl_2_ as reported previously.^31^ The assembled DNA channel and baseplate were then mixed at an equimolar ratio and incubated at 37°C for 48 hours for dimerization. The complete NanoTrap was purified using rate-zonal centrifugation^33^ through a 15–45% glycerol gradient in 1×TE + 10 mM MgCl_2_ in an SW 55 rotor (Beckman Coulter). Fractions were collected after a 1 hr centrifugation at 50 k rpm. Typically, 5 μL of each fraction was loaded in a 1.5% agarose gel (0.5× TBE, 10 mM MgCl_2_) with 0.5 μg/mL ethidium bromide (EtBr). A 1 kb DNA ladder (New England Biolabs) was run in parallel with samples. Electrophoresis was carried out at 5 V/cm for 120 min in 0.5× TBE, 10 mM MgCl_2_. Gels were imaged on a Typhoon FLA 9500 scanner (GE Healthcare). Fractions containing desired DNA structures (determined by agarose gel electrophoresis) were collected, and the buffer was changed to 1×TE + 10 mM MgCl_2_ using Amicon Ultra centrifugal filters with 100 kD cutoff (EMD Millipore). The purified NanoTraps were then stored at −20°C.

### Attaching FG-nups to DNA NanoTrap

DNA-conjugated nups was added to DNA NanoTraps at 1.5× excess over the number of handles (e.g., 5 nM NanoTrap × 48 handles × 1.5= 360 nM FG-nup-DNA) in 1 ×TE buffer with 15 mM MgCl_2_. The mixture was kept at 37°C for 2 hr to allow handle-to-anti-handle hybridization. The products were purified by rate-zonal centrifugation, as described previously,^31^ through a 15–45% glycerol gradient in the hybridization buffer (1×TE buffer with 15 mM MgCl_2_).

### Penetration assay

#### Sample preparation

We tested the diffusion barriers formed by FG-nups by incubating the FG-nup-gated NanoTraps with a series of fluorescently labeled molecules (reporters) that ranged from 7 kD to 106 kD: an Alexa488-prey (7 kD), a GFP-SNAP-prey (53 kD), and an MBP-GFP-SNAP-prey (106 kD). Briefly, 3 nM NanoTraps containing various FG-nup configurations were incubated in separate test tubes with 1 μM reporters of different sizes for 1.5 hours at 37°C. Empty NanoTrap was incubated with the same set of reporters under identical conditions.

#### SDS-Agarose gel electrophoresis

Samples were loaded in a SDS-agarose gel (1.5% agarose in 0.5×TBE, 10 mM MgCl_2_, and 0.05% SDS). Electrophoresis was carried out at 5.8 V/cm for 90 min in 0.5×TBE buffer containing 10 mM MgCl_2_ and 0.05% SDS. Gels were imaged on a Typhoon FLA 9500 scanner (GE Healthcare) for the in-gel fluorescence (GFP or Alexa Fluor 488) first, stained with ethidium bromide (EtBr), and then imaged again for the EtBr fluorescence. For EtBr staining, the gel was first soaked in deionized H2O and shaken for 1 hr to remove SDS, and then submerged in an EtBr solution (Sigma-Aldrich, 20,000× dilution in H2O to 0.5 μg/mL) for 1 hr. Gels were destained for 1 hour in deionized H2O before imaging.

#### Image analysis

The gel images were analyzed using ImageJ (v2.1.0) using the built-in gel analyzing tool for the band intensities. To account for possible concentration variation among the NanoTrap samples, all NanoTrap bands’ GFP/Alexa Fluor 488 fluorescence (from the trapped reporter molecules) were normalized against their EtBr fluorescence. The normalized fluorescence of the empty NanoTrap was set as a reference with 100% penetration; the penetration of a certain reporter through an FG-nup-gated NanoTrap was quantified by dividing the normalized fluorescence of the NanoTrap band by that of the reference band and expressed as percentages (**Figure 3** and **4**).

### Negative-Stain Transmission Electron Microscopy

Negative-stain TEM was used to visualize the DNA channel and baseplate, as well as empty and FG-nup-gated NanoTraps. Typically, samples (5 μL) were loaded onto a glow discharged Formvar/carbon-coated copper grid (400 mesh, Electron Microscopy Sciences) and stained with 2% uranyl formate. Imaging was performed on a JEOL JEM-1400 Plus microscope operated at 80 kV with a bottom-mount 4k×3k CCD camera (Advanced Microscopy Technologies).

### Attaching AuNP to DNA NanoTrap

Thiol-labeled prey-oligo (41 μM) was mixed with phosphine-treated 5 nm AuNP (200 nM, Ted Pella) in 50 mM NaCl, 1× TBE buffer (44.5 mM Tris, 44.5 mM boric acid, 1 mM EDTA).^49^ The mixture was covered with aluminum foil and agitated in a ThermoMixer (Eppendorf) under r.t. at 300 rpm for ~40 hr. Subsequently, the DNA-conjugated AuNP was purified and washed with 0.5× TBE buffer using Amicon Ultra centrifugal filters with 50 kD cutoff (EMD Millipore). To characterize the product, 5 μL of AuNP was loaded in a 3% agarose gel, which was run in 1× TAE buffer (40 mM Tris, 20 mM acetic acid, 2 mM EDTA) at 10 V/cm for 30 mins (**Figure S4**). OD_520_ of the resuspended AuNPs was measured to determine the AuNP concentration. The purified DNA-conjugated AuNPs were stored at 4 °C until use.

For AuNP attachment, empty NanoTrap (2 nM) was incubated with prey-oligo-conjugated AuNP (2 nM) for 1.5 hours at 37°C. The mixture was imaged by negative-stain EM to visualize the immobilization of AuNPs inside the NanoTraps.

### Statistical analysis

The data analysis was performed using the SPSS 26.0 software package (IBM, United States). Unless noted otherwise, all statistical data were expressed in mean ± standard error of the mean (SEM). Two-tailed t-tests were applied to evaluate the differences between top and bottom arranged nucleoporins, two-way ANOVA and Tukey’s multiple comparisons test was applied to evaluate the difference between mixed-nup NanoTraps and Nup100-only NanoTraps. Detailed statistics data were shown in **Table S1**. P < 0.05 were considered statistically significant.

## SUPPLEMENTAL INFORMATION

Figures S1–S8; Table S1.

## ACKNOWLEDGMENTS

We thank the Lin Lab for discussion; This work was supported by National Institutes of Health grants R01 GM132114 and P50 AI150481 (C.L.), R01 GM105672 (C.P.L.), and R21 GM109466 (C.L., C.P.L. and T.J.M.), as well as a Singapore Agency for Science, Technology and Research Graduate Scholarship (Q.X.).

## AUTHOR CONTRIBUTIONS

Q.S., P.D.E.F., T.M., C.P.L., and C.L. conceived and designed research; Q.S., T.T., Q.X. and P.D.E.F. performed the experiments; Q.S., T.T., Q.X. analyzed data; Q.S., T.T., Y.X., C.P.L., and C.L. interpreted the data. Q.S. T.T., Q.X. wrote the original draft; and Q.S., T.T., Q.X., C.P.L., and C.L. reviewed and edited the manuscript. All authors participated in the discussions.

