## Supplementary Information for "A DNA-origami NanoTrap for studying the diffusion barriers formed by Phe-Gly-rich nucleoporins"

Supporting Figures S1–S8, Table S1

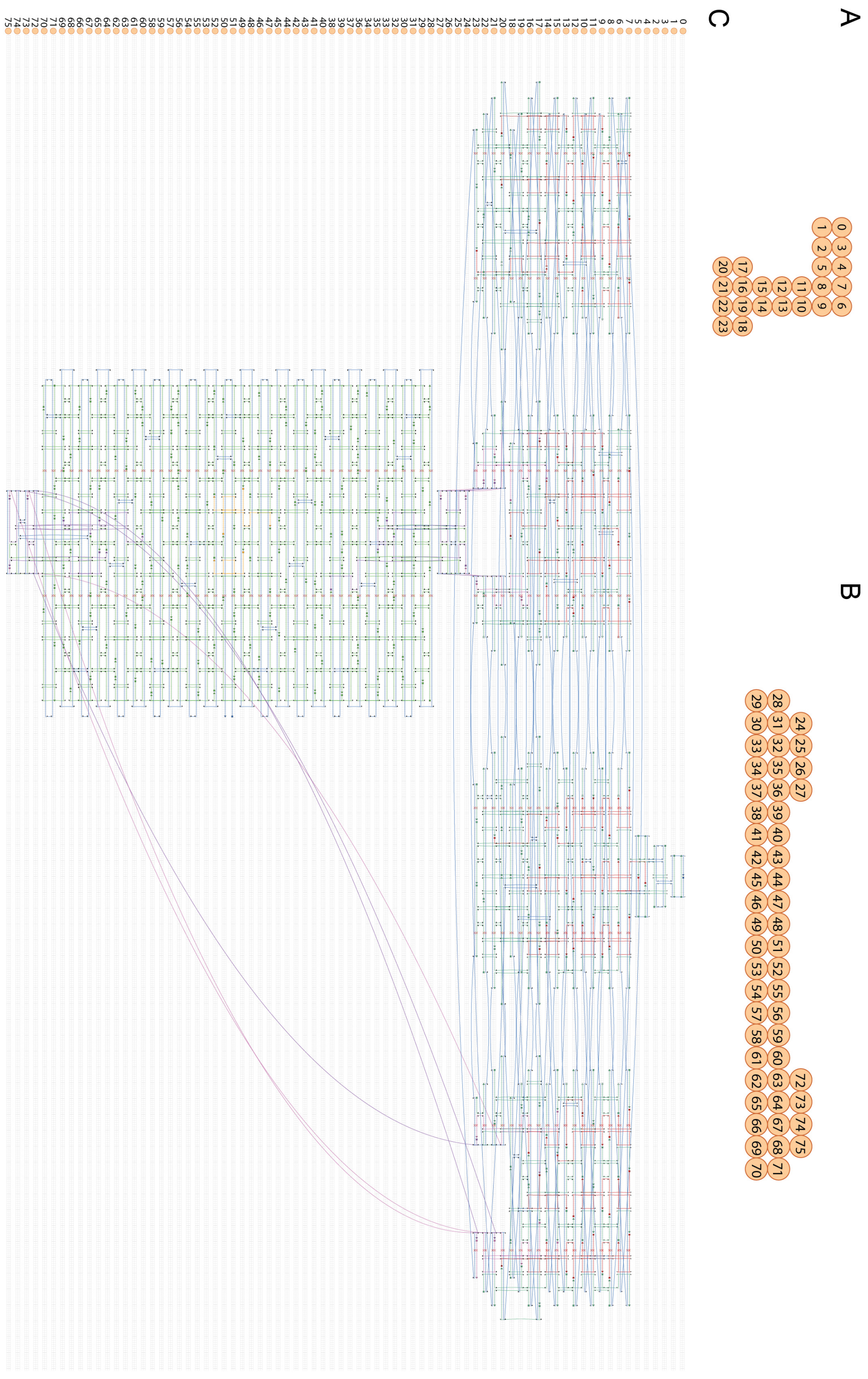


**Figure S1**. **DNA-origami designs rendered in caDNAno.**

(A) Cross-section view of the channel; (B) Crosssection view of the baseplate; (C) strand diagrams for the NanoTrap design. The scaffold strand is in blue. Staple strands for inner handles, teeth, and bait are shown in red, purple, and orange, respectively.

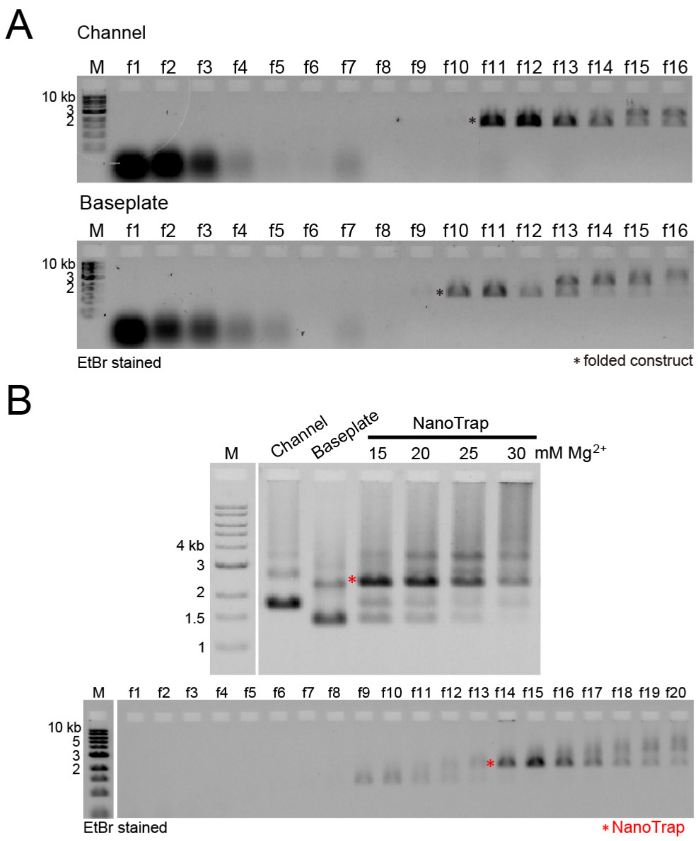


**Figure S2. DNA-origami assembly and purification.**

(A) Folded DNA channel and baseplate purified by rate-zonal centrifugation. Agarose gel (1.5%) electrophoreses show the enrichment of DNA channel (top) and baseplate (bottom) in fractions 11–13 and 10–12, respectively. Well-folded nanostructure bands are denoted by an asterisk; (B) Agarose gel electrophoreses show the channel and baseplate dimerization yield at different $MgCl_2$ concentrations (top) and the enrichment of NanoTrap in fractions 14–16 after rate-zonal centrifugation (bottom). NanoTrap bands are denoted by an asterisk.

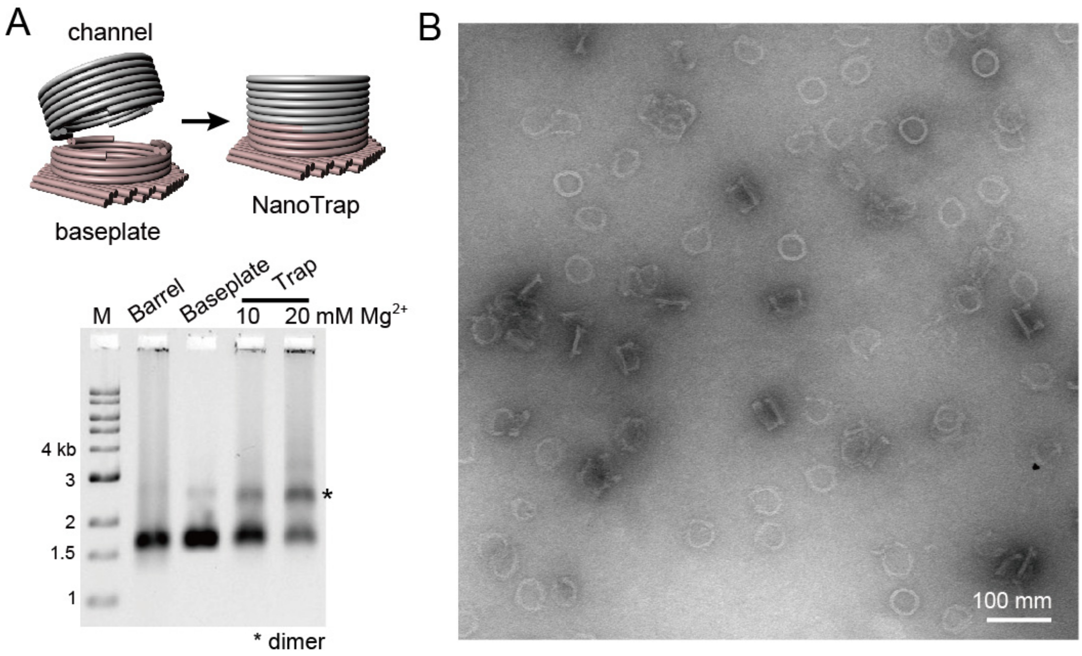


**Figure S3. DNA NanoTrap with a ring-shaped channel**

(A) Cartoon models of the ring-shaped NanoTrap assembly (top) and the analysis of dimerization yields by agarose gel electrophoresis (bottom); (B) Negative-stain TEM image of the ring-shaped NanoTrap. Scale bar: 100 nm. Note the inferior assembly efficiency compared to the NanoTrap used in this study (**Figure 1** and **S2**).

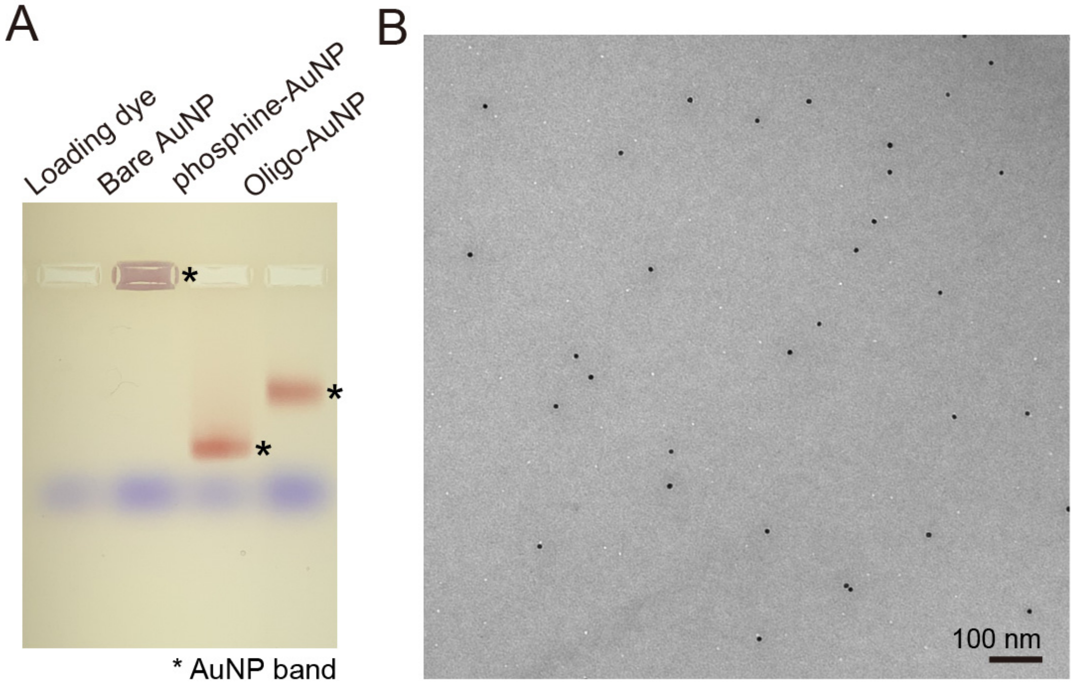


**Figure S4. Preparation of prey-oligo-conjugated AuNP**

(A) Agarose electrophoresis showing the different mobilities of bare AuNP (stuck in the well), phosphine-treated AuNP, and prey-oligo-conjugated AuNP. AuNP bands are denoted by an asterisk. (B) A TEM image of the prey-oligo-conjugated 5 nm AuNPs with no signs of aggregation. Scale bar: 100 nm.

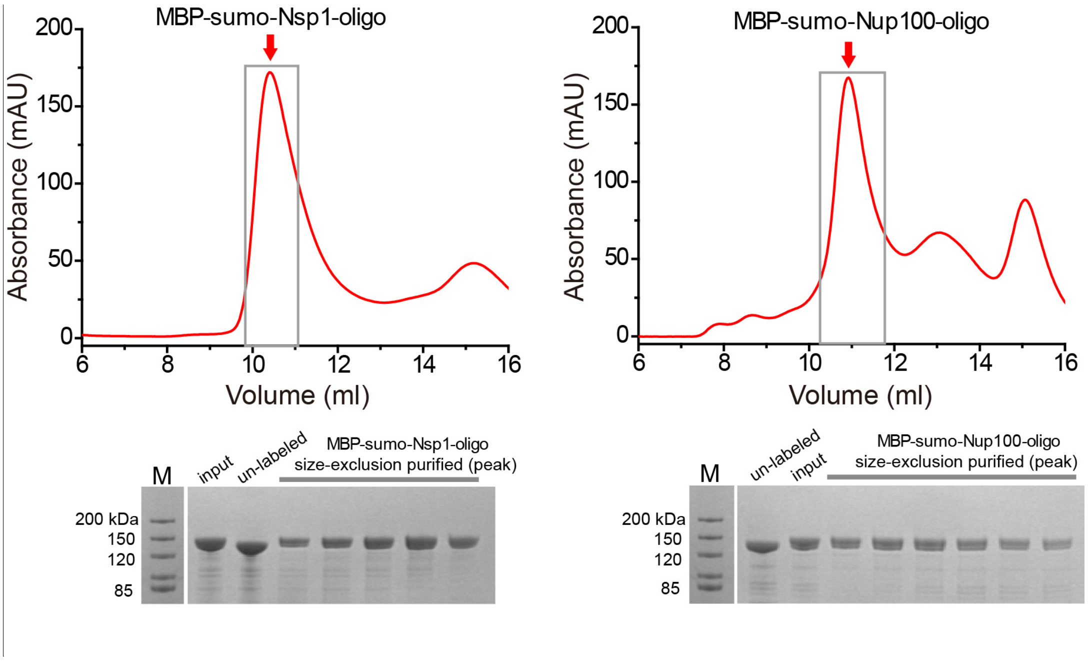


**Figure S5. MBP-sumo-Nsp1-oligo and MBP-sumo-Nup100-oligo purification.** The DNA-conjugated Nsp1 and Nup100 are marked by a red arrow in their respective size exclusion chromatography graphs. SDS-PAGE show the fractions containing purified nup-DNA conjugates (denoted by a gray box in the chromatography graphs).

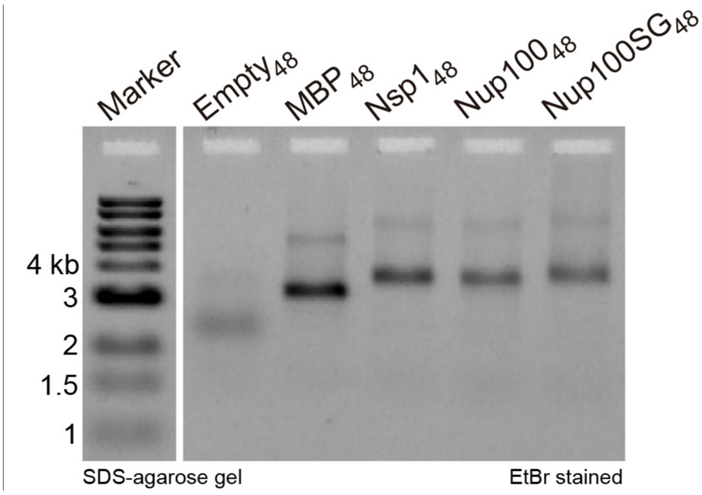


**Figure S6. NanoTraps characterized by SDS-agarose gel electrophoresis.**

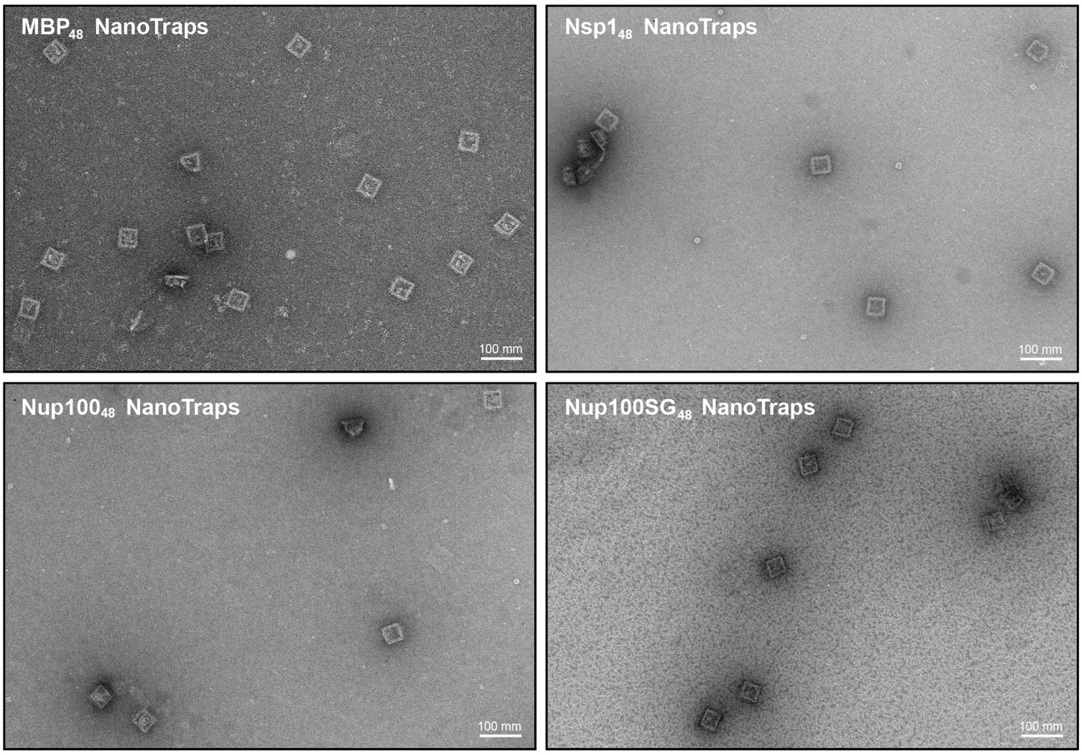


**Figure S7. Negative-stain EM images of various protein-gated NanoTraps.**

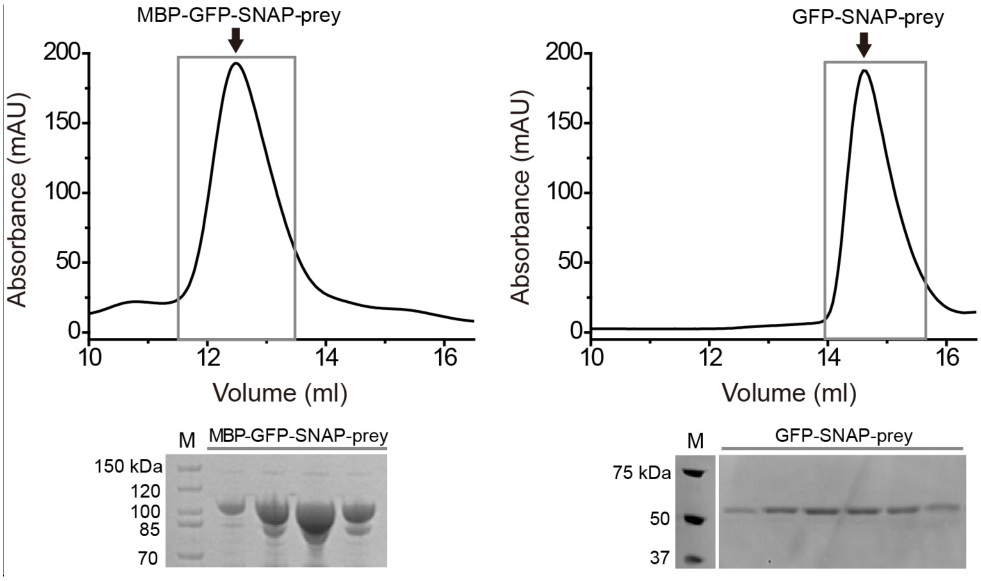


**Figure S8. MBP-GFP-SNAP-prey and GFP-SNAP-prey purification.** The DNA-conjugated MBP-GFP-SNAP and GFP-SNAP are marked by an arrow in their respective size exclusion chromatography graphs. SDS-PAGE show the fractions containing purified protein-DNA conjugates (denoted by a gray box in the chromatography graphs).

**Table S1. Data of statistical analysis**

| Statistics data of t-test (Nup100 located at top and bottom of the NanoTrap) | | | | |
|---|---|---|---|---|
| | F-test (F, DFn, Dfd) | P-value of F-test | t-test (t, df) | P-value |
| 12 top vs 12 bottom | 2.337, 2, 2 | 0.5993 | 7.821, 4 | 0.0014 |
| 24 top vs 24 bottom | 2.252, 2, 2 | 0.615 | 2.711, 4 | 0.0535 |
| 36 top vs 36 bottom | 1.286, 2, 2 | 0.875 | 1.744, 4 | 0.1562 |

| Statistics data of t-test (Nsp1 located at top and bottom of the NanoTrap) | | | | |
|---|---|---|---|---|
| | F-test (F, DFn, Dfd) | P-value of F-test | t-test (t, df) | P-value |
| 12 top vs 12 bottom | 3.910, 2, 2 | 0.4073 | 0.5298, 4 | 0.6243 |
| 24 top vs 24 bottom | 1.912, 2, 2 | 0.6869 | 0.2244, 4 | 0.8335 |
| 36 top vs 36 bottom | 1.981, 2, 2 | 0.6709 | 2.424, 4 | 0.0725 |

| Statistics data of Tukey's multiple comparisons test (mixed vs Nup100) | | | | | |
|---|---|---|---|---|---|
| Test details | Mean 1, mean2, mean diff. | 95.00% CI and SE of diff. | N1, N2 | Q-value, df | P-value |
| $Empty_{36}Nup100_{12}$ | 15.63, 33.98, -18.35 | -35.24 to -1.457, 6.91 | 3, 3 | 3.755, 36 | 0.0308 |
| $Nup100_{12}Empty_{36}$ | 40.62, 71.39, -30.77 | -47.66 to -13.88, 6.91 | 3, 3 | 6.296, 36 | 0.0002 |
| $Empty_{24}Nup100_{24}$ | 15.34, 17.16, -1.821 | -18.71 to 15.07, 6.91 | 3, 3 | 0.3727, 36 | 0.9625 |
| $Nup100_{24}Empty_{24}$ | 29.41, 25.14, 4.265 | -12.63 to 21.16, 6.91 | 3, 3 | 0.8728, 36 | 0.8117 |
| $Empty_{12}Nup100_{36}$ | 19.8, 14.6, 5.2 | -11.69 to 22.09, 6.91 | 3, 3 | 1.064, 36 | 0.7341 |
| $Nup100_{36}Empty_{12}$ | 24.23, 22.37, 1.853 | -15.04 to 18.74, 6.91 | 3, 3 | 0.3791, 36 | 0.9612 |
